# Accurate quantitation of 16S gene copies in low biomass samples post-antibiotic treatment through deep sequencing with a balanced nucleotide synthetic spike-in approach

**DOI:** 10.1101/2024.09.27.615519

**Authors:** Chia-Chi Chang, Licai Huang, Yimei Jin, Mohamed A. Jamal, Eiko Hayase, Taylor Halsey, Miriam R. Ortega, Dung Pham, Israel Glover, Ivonne Flores, Lauren McDaniel, Rishika Prasad, Christine Peterson, Robert R. Jenq

## Abstract

The microbiota significantly impacts health and treatment outcomes. While 16S rRNA gene sequencing reveals relative bacterial abundances, it does not provide absolute quantification. We developed a cost-effective solution incorporating synthetic DNA standards designed to ensure balanced nucleotide representation at each position. These standards are spiked into samples before DNA extraction, enabling simultaneous quantification of both relative and absolute bacterial abundances. We applied this method to samples collected from mice and patients, both before and after antibiotic treatment. Our approach showed a reduction in total bacterial density in mice and patients post-antibiotic treatment. This spike-in standard method can be adapted to samples with varying bacterial densities, allowing quantification of absolute taxonomical abundances without the need for an additional quantitative PCR assessment. Our approach also improves sequencing quality scores for low biomass samples.

## Background

Advancements in next generation sequencing technologies have facilitated a greater understanding of interactions between microbial communities and the hosts that harbor them.^1^ However, interpretation of assays quantifying microbial communities that solely rely on relative abundances can be misleading in the absence of concomitant absolute quantification, especially for low bacterial biomass specimens, including samples collected in settings of antibiotic treatment. In one example, quantitative PCR (qPCR) revealed that pancreatic tumors from KPPC (LSL-Kras^G12D/+^; Trp53^loxP/loxP^; Pdx1-Cre) mice showed higher bacterial densities compared to normal pancreases from healthy littermate mice. qPCR can also be used to assess bacterial density after antibiotic treatment. Antibiotic treatment can sometimes substantially deplete the gut and systemic microbiota, depending on the antibiotic spectrum of activity, bioavailability, and sensitivity of commensal organisms. Treatment with an antibiotic cocktail led to altered tumor immune profiling in KPPC;Col1^pdxKO^ mice, which possess a genetic deletion of Col1 in cancer cells of KPPC mice, and resulted in shortened survival in KPPC;Col1^pdxKO^ mice compared to those without antibiotic treatment.^2^ In another study, patients with osteoporosis were found to have an increase in the absolute abundances of Bacteroidetes phylum and *Bacteroides* genus in comparison with the controls.^3^ Similarly, a study on the role of the microbiome in osteoarthritis found that greater relative and absolute levels of intestinal *Streptococcus* species were correlated with increased osteoarthritis-related knee pain.^4^ In a study involving patients with Crohn’s disease, investigators used a method that combined counting microbial cells into the sequencing workflow. This approach allowed them to measure the quantity of microbes more accurately. They discovered that the microbial loads varied greatly among individuals. The key finding was that the changes in the microbiome were primarily due to an overall reduction in the number of microbial cells, rather than an increase in specific disease-related microbes.^5^ Moreover, in a recent study, patients with higher *Staphylococcus aureus* cell numbers and total bacterial load in lesion samples were significantly more likely to have severe atopic dermatitis compared to patients with mild or moderate atopic dermatitis.^6^ Thus, absolute quantification of bacterial densities, in addition to quantifying relative contributions of bacterial subpopulations, can be critical for developing an accurate understanding of whether bacterial populations are expanding or shrinking over time and their associations with disease in patients and preclinical models.^7^

qPCR techniques allow precise determination of absolute abundances of target DNA sequences and can be used to quantify microbial density, but are not always routinely performed in the setting of next-generation sequencing (NGS).^8^ Performing qPCR requires adequate DNA yields and can sometimes be challenging with low biomass samples including samples collected after antibiotic treatment.^7^ An elegant, alternative approach involves introducing known amounts of spike-in references prior to NGS, which can enable streamlined simultaneous assessment of both absolute and relative abundances for potentially richer interpretation of microbiota samples. In the past few years, several studies have reported applying spike-in quantitative methods for microbiome applications, including synthetic spike-in markers or unusual and distinguishable microorganisms.^9-11^ Tourlousse et al. designed synthetic spike-in standards *in silico* as an internal standard, allowing robust sample tracking^12^ and detection of cross-contamination in microbiome community analysis by 16S rRNA gene amplicon sequencing. Spiking a known concentration of exogenous bacteria is another strategy that can be used to quantify total absolute bacterial abundances.^10^ A novel NGS approach that combines viability reagent and spike-in control steps can discriminate between viable and dead microbial cells and accurately estimate quantitative microbiota analysis in food.^6^ Similar strategies have been employed for different NGS approach such as shotgun metagenomic sequencing.^13^ Certain principles have been identified to be important for such methods. First, the introduced references should either be ideally absent or at a minimum rarely present in actual samples. Next, the amount of spike-in internal references needs to be carefully optimized, striking a balance between not overloading and reducing the reads generated from the actual microbial DNA, but being abundant enough to allow accurate quantification. Third, an appropriate normalization method is important for generating a standard curve for absolute quantification.

In this study, we identified that an additional factor, balanced nucleotide representation of the spike-in DNA, can be important for successful performance of this approach. This led us to develop a nucleotide-balanced four spike-in approach called “balanced synthetic spike-in,” or BSSI. When used for 16S rRNA sequencing, we found that BSSI allows simultaneous quantification of total absolute microbial abundance and relative abundances without the need for performing an additional 16S qPCR. A mixture of four synthetic spike-in plasmids were added directly into lysis buffer, co-extracted and PCR-amplified, enabling calculation of absolute abundances of prokaryotic 16S rRNA copies. Importantly, the BSSI approach significantly improves sequencing quality scores, compared to an imbalanced commercially-available spike-in method. Additionally, we found that this approach performed well across a considerable dynamic range, allowing quantification of total bacterial densities across five orders of magnitude. It also performed more robustly than 16S rRNA qPCR for analyzing very low-density fecal samples collected from antibiotic-treated mice and patients. We believe this strategy can potentially be applied to a variety of research samples, allowing quantification of absolute abundances of prokaryotic 16S rRNA copies and determination of bacterial composition in a single deep sequencing assay.

## Methods

### Design of synthetic 16S spike-in mix standards and plasmid construction

The design of a mixture of balanced DNA standards includes four synthetic spike-ins (SP1, SP2, SP3, and SP4), each with sequences containing 253 nucleotides with variable GC content (57.3, 52.6, 47.4 and 42.7% of GC). In each of the four DNA sequences, at every base position, there are the four complementary bases: A, T, C, and G. Constant adapter sequences were added to both the 5’ and 3’ ends of all synthetic DNA spike-ins, which results in a total length of 292 base pairs, as listed in Table 1. The 5’ end constant sequence is ‘5’-gtgccagcagccgcggtaa,’ and the 3’ end constant sequence is ‘5’-attagataccctggtagtcc,’ which are used for mimicking amplification of the 16S fourth hypervariable (V4) region. The four synthetic DNA standards were cloned into pUC57-Kan plasmids using the EcoRV digestion site to generate spike-in DNA plasmids, SP1-pUC57-Kan, SP2-pUC57-Kan, SP3-pUC57-Kan, and SP4-pUC57-Kan. These constructs were synthesized and purchased from GenScript Biotech. To produce the plasmids, *Escherichia coli* DH5α (Thermo Fisher Scientific Cat. EC0112) transformants were grown on LB medium supplemented with kanamycin (50 μg/mL) for selection. To create a mixture of imbalanced DNA standards, four synthetic spike-ins (SP1, SP2, SP5, and SP6), each with a variable sequence, were synthesized by GenScript Biotech. The sequences of SP5 and SP6 are listed in Table 2. They were cloned into a pCR4-Blunt II-TOPO vector to generate SP1-pCR, SP2-pCR, SP5-pCR, and SP6-pCR plasmids, using the Zero Blunt PCR Cloning Kit (Thermo Fisher Scientific, Cat. K280020), following the manufacturer’s protocol.

### Patient microbiome analyses

We identified four patients diagnosed with acute myeloid leukemia (AML) or myelodysplastic syndrome (MDS) who underwent hematopoietic stem cell transplantation in 2023 and received treatment with the antibiotic cefepime (Table 3). Stool samples were collected from the patients before and during the cefepime treatment. Samples designated as collected during cefepime treatment were collected four days to seven days after beginning continuous cefepime treatment, typically administered intravenously every 8 hours. Following collection, the stool specimens were initially preserved at 4°C for a period ranging from 24 to 48 hours, and subsequently portioned into aliquots for long-term storage at −80°C. Informed consent was obtained from all the study participants, and the study was approved by the Institutional Review Board of the University of Texas MD Anderson Cancer Center under protocol number 2021-0206.

### Animal experiment

The IACUC committee approved the care and use of laboratory animals in our study prior to the initiation of the experiments. Commercial chow and water were autoclaved or irradiated before use. The C57BL/6J mice, acquired from The Jackson Laboratory, were housed in a barrier facility with a 12-hour light-dark cycle. After an acclimation period, 10-week-old C57BL/6J mice were given an antibiotic cocktail in drinking water (vancomycin - 1000 mg/L, ampicillin - 1000 mg/L, enrofloxacin - 250 mg/L) for 4 days and then were off antibiotics for 7 days. Stools were collected before, on the fourth day of antibiotic treatment, and seven days after discontinuing the antibiotic water. The CD11c-Cre+ Stat3f/f mice (Stat3Δ/Δ)^14^, obtained from Dr. Stephanie Watowich’s laboratory, were generated by breeding B6.Cg-Tg (Itgax-cre)1-1Reiz/J mice (CD11c-Cre; 008068; The Jackson Laboratory) with Stat3f/f mice^15,16^. Five-week-old CD11c-Cre+ Stat3f/f mice (Stat3Δ/Δ) were treated with an antibiotic cocktail in drinking water (vancomycin - 1000 mg/L, ampicillin - 1000 mg/L, enrofloxacin - 250 mg/L) for five days, and fecal pellets were collected on the fifth day of antibiotic treatment.

### Genomic DNA extraction with synthetic spike-in standards

Fecal samples were collected from patients and mice and were weighed prior to DNA extraction. *Escherichia coli* DH5α cells (Thermo Fisher Scientific, Cat. EC0112) were counted and prepared in 10^6^, 10^7^, 10^8^, 10^9^ and 10^10^ cells in the tubes for DNA isolation. Bacterial cell counts and viability were determined by staining with Syto BC dye (Thermo Fisher Scientific, Cat. S34855) and propidium iodide (Nexcelom Bioscience, Cat. CS1-0109) and assessed using an automated cell counter (Nexcelom Bioscience). The viability is above 98%. The spike-in DNA plasmid mixture of SP1-pUC57-Kan, SP2-pUC57-Kan, SP3-pUC57-Kan, and SP4-pUC57-Kan were mixed in an approximate 1:1:1:1 equimolar ratio. A total of 0.5 ng of the standards mixture was pre-mixed with 600 μl of InhibitEx lysis buffer (Qiagen, Cat. 19593) and then added to bacterial cell pellets or fecal samples for DNA extraction, followed by an intensive bead-beating step using metal beads (Qiagen, Cat. 6999) and zirconia beads (BioSpec, Cat. 11079101Z). Bacterial genomic DNA was isolated using the QIAamp fast DNA stool kit (Qiagen, Cat. 51604), in accordance with the manufacturer’s protocol.^17^ For the imbalanced sequencing diversity assay, the *Akkermansia muciniphila* (MDA-JAX AM001)^18^ and *Bacteroides thetaiotaomicron* (MDA-JAX BT001)^19^ isolates were mixed in an approximate 1:1 ratio based on cell count and spiked with an imbalanced plasmid mixture consisting of SP1-pCR, SP2-pCR, SP5-pCR, and SP6-pCR plasmids before the genomic DNA extraction. Both MDA-JAX AM001 and MDA-JAX BT001 isolates were murine-derived and recovered from the feces of C57BL/6 female mice, were verified by Sanger sequencing and MALDI Biotyper (Bruker) before use.

### 16S rRNA V4 sequencing and microbiome data analysis on spike-in standards curve

The V4 region of the 16S rRNA gene was amplified by PCR from each extracted and purified genomic DNA using the 515 forward (5’-GTGYCAGCMGCCGCGGTAA) and 806 reverse (5’-GGACTACNVGGGTWTCTAAT) primer pairs. The amplicon pools were purified with a QIAquick gel extraction kit (Qiagen, Cat. 28704) and sequenced on the Illumina MiSeq sequencer using a 2 × 250 bp paired-end protocol. The experiments assessing the impact of different spike-in standards on sequencing run quality used a 2 × 150 bp paired-end protocol. The reads were merged, dereplicated, and length-filtered utilizing VSEARCH.^20^ Following denoising and chimera calling using the UNOISE3 commands,^21^ the unique sequences, or zero-radius OTUs (ZOTUs), were taxonomically classified using Mothur.^22^ with the SILVA database version 138. Alpha and beta diversity metrics were generated in QIIME 2.^23^ The Q scores, pass filter rates, percentage of PhiX alignment and cluster density were generated by sequencing analysis viewer software (Illumina). The abundances of spike-in standards SP1, SP2, SP3, and SP4 sequences were assigned to artificial taxonomical groups and their relative abundance was calculated alongside other taxonomical groups. The spike-in standard formula was generated by adding to 0.5 ng of spike-in DNA mixture with seven different amounts (500 ng, 50 ng, 5 ng, 0.5 ng, 0.05 ng, 0.005 ng, and 0.0005 ng) of pCR4-*Blautia* plasmid, which contains full-length 16S rRNA gene sequences cloned from *Blautia luti DSM 14534* species with 95.29% identity (accession NR_114315.1) ^24^.The full-length sequence of the 16S *Blautia* species is listed in Table 1.

The formula for the spike-in standard curve was calculated using the known 16S gene copies of the pCR4-*Blautia* plasmid (in log_10_) as the x-axis, and the corresponding points of abundance of the pCR4-*Blautia* plasmid, calculated through different normalization methods- logit, arscine and anscombe-as the y-axis. The formula for logit transformation is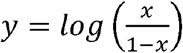. The equation for arcsine normalization is 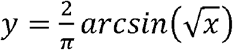. The anscombe normalization is represented by the equation 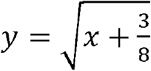 The formula y = 1.7745x - 13.184 was used to describe the relationship between logit-transformed relative abundances and log-transformed absolute bacterial densities. (Fig. S1A, S1B and S1C) The standard curve has an R-squared value >0.99, as shown in Table 4. Other taxonomic classifications assigned at the species, genus, family, class, order, and phylum levels were ranked in descending order of variance in abundance.

### 16S rRNA quantitative PCR

Six pCR4 plasmid standards, representing a 10-fold dilution series (200 pM, 20 pM, 2 pM, 0.2 pM, 0.02 pM, and 0.002 pM), were used to generate the standard curve for 16S rRNA real-time qPCR for mouse and human samples. *Escherichia coli* DNA (Sigma Cat. MBD0013) was used to generate a standard curve for the *Escherichia coli* DH5α bacterial cell experiment. The qPCR assay was prepared with Kapa Fast SYBR Green Master Mix (Kapa Biosystems, Cat. KK4824) and run on the QuantStudio 6 Flex Real-Time PCR System (Applied Biosystems) using the 926-forward primer (5’-AAACTCAAAKGAATTGACGG) and 1062-reverse primer (5’-CTCACRRCACGAGCTGAC) pairs. The PCR conditions were 95°C for 10 minutes, followed by 40 cycles of 95°C for 15 seconds and 60°C for 1 minute. A melting curve analysis was performed after amplification.^25^

### Statistical Analysis

All data were analyzed using GraphPad Prism version 10.0.3 and R Studio version 2023.09.1+494. Student’s t-tests, Pearson’s correlation and proportion test were used when appropriate. Mann-Whitney U tests were used to compare data between two groups when the data did not follow a normal distribution. GraphPad Software P values < 0.05 were considered significant.

## Results

### Choice of synthetic 16S spike-in mixture standards

All four balanced synthetic 16S sequence constructs were 253 base pairs in length and designed *in silico* to align poorly to all 16S rRNA sequences in the NCBI 16S ribosomal microbial database (Table 1). Additionally, representation by the four individual nucleotide bases (A, T, C, and G) was purposefully balanced across the four constructs at each nucleotide position (Fig. 1A). These four sequences, SP1, SP2, SP3 and SP4, were then flanked by constant adapter sequences to allow amplification of these constructs using Earth Microbiome V4 region-specific primers.^26-28^ We first evaluated the ability of the 515F and 806R primer pairs to amplify the spike-in standards. Agarose gel revealed that all four spike-in standards can be successfully amplified by the 16S V4 515F - 806R primer pair (Fig. S2). ^26,27^ The GC content of all spike-ins standards ranged from 42.7% to 57.3% of nucleotides. The full-length spike-in references were synthesized and cloned into the pUC57-Kan plasmids and verified by Sanger sequencing. The spike-in standards were pre-mixed in a 1:1:1:1 equimolar ratio and stored at - 30°C before use. The standards mixture was then added to lysis buffer during the DNA lysis step followed by standard DNA extraction, library preparation and 16S sequencing (Fig. 1B).

**Figure 1.**
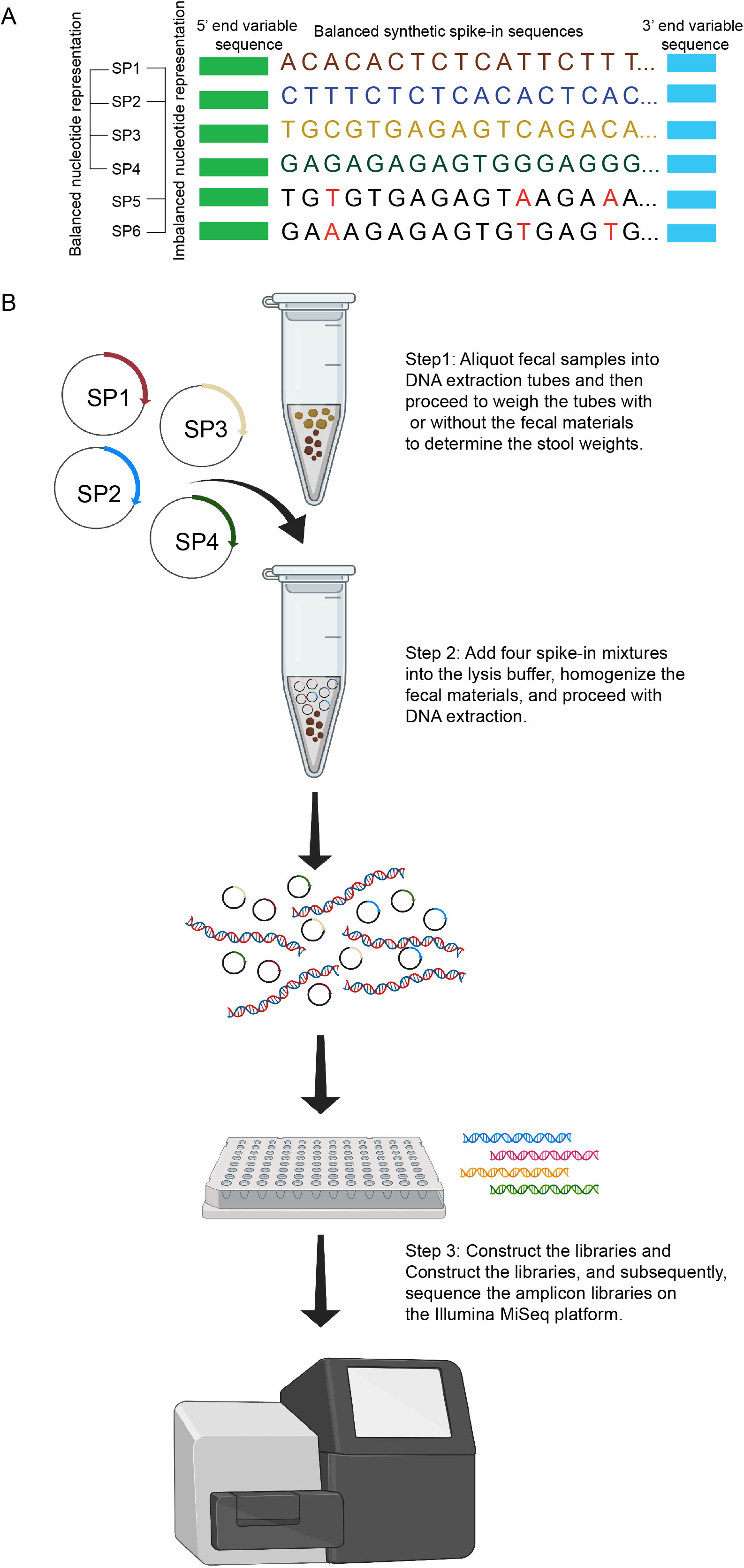
Synthetic spike-in amplicon sequences and 16S spike-in workflow. (A) In the sequence of four synthetic sequences, SP1, SP2, SP3, and SP4, nucleotide bases (A, T, C, and G) were purposefully balanced across the four constructs at each nucleotide position. The diagram also includes the design of imbalanced nucleotides, SP5 and SP6, which have imbalanced nucleotide bases highlighted in red when mixed with the SP1 and SP2 standards. (B) The 16S spike-in standards workflow: Four synthetic spike-in DNAs, pre-mixed in an equal molar ratio, were added during the DNA lysis step, followed by 16S PCR amplification and next-generation sequencing.

### Characterization of the spike-in standards for absolute quantification

To assess the accuracy of our spike-in method, we cultured *Escherichia coli* DH5α, a commercially available strain, and prepared samples consisting of five different cell densities: 10^6^, 10^7^, 10^8^, 10^9^ and 10^10^ bacterial cells. BSSI standards were added to each of these *Escherichia coli* samples before proceeding with cell lysis, followed by DNA isolation, 16S library preparation, and deep sequencing. The 16S rRNA gene copies for each these samples were determined after generating a spike-in standard curve with additional samples including varying quantities of a plasmid encoding a single 16S rRNA gene copy (Table 4 and Fig. S1). To evaluate the performance of spike-in standard curve for absolute quantification across a range of 16S gene densities, we assessed the R-squared values of standard curves generated by plotting the normalized abundance of pCR4-*Blautia* plasmid gene copy numbers. We compared using logit, arcsine, and anscombe normalization methods on the y-axis against the log_10_-transformed known 16S gene copies of the pCR4-*Blautia* plasmid on the x-axis. Logit normalization was superior to arcsine and anscombe normalization methods, with a higher R-squared value (R^2^ = 0.9972) (Fig. S1A) compared to arcsine (R^2^ = 0.9617) (Fig. S1B) and anscombe (R^2^ = 0.9229) (Fig. S1C) normalization methods, without flattening of the linear relationship at the extremes of tested densities. We then performed absolute quantification using the spike-in standard curve with logit normalization for the abundance of pCR4-*Blautia* plasmid gene copy numbers on samples with various *Escherichia coli* DH5α cell densities. In the samples containing 10^6^, 10^7^, 10^8^, 10^9^, and 10^10^ bacterial cells, the mean 16S rRNA gene copies were 2.49E+06, 2.54E+07, 1.75E+08, 1.85E+09, and 2.25E+10, respectively (Fig. 2A left panel). DNA samples extracted from the five different bacterial count samples, as described above, were also subjected to traditional 16S qPCR to evaluate total absolute abundances. The qPCR primers were designed to amplify the hypervariable regions of V6-V7, whereas the deep sequencing method used primers to amplify only the V4 region. This analysis utilized commercially available *Escherichia coli* genomic DNA to generate the standards. Pearson correlation analysis demonstrated a high degree of correlation (r = 0.99, P < 0.001) between the results obtained from the spike-in standard method and the 16S rRNA qPCR method (Fig. 2A right panel). These findings suggest that spike-in standards can be co-amplified with bacterial genomic DNA using 16S V4 primer pairs and provide a reliable method for quantifying total absolute abundances through deep sequencing. Additionally, the spike-in standard approach performed well with a large dynamic range, allowing quantification of total bacterial densities across five orders of magnitude.

**Figure 2.**
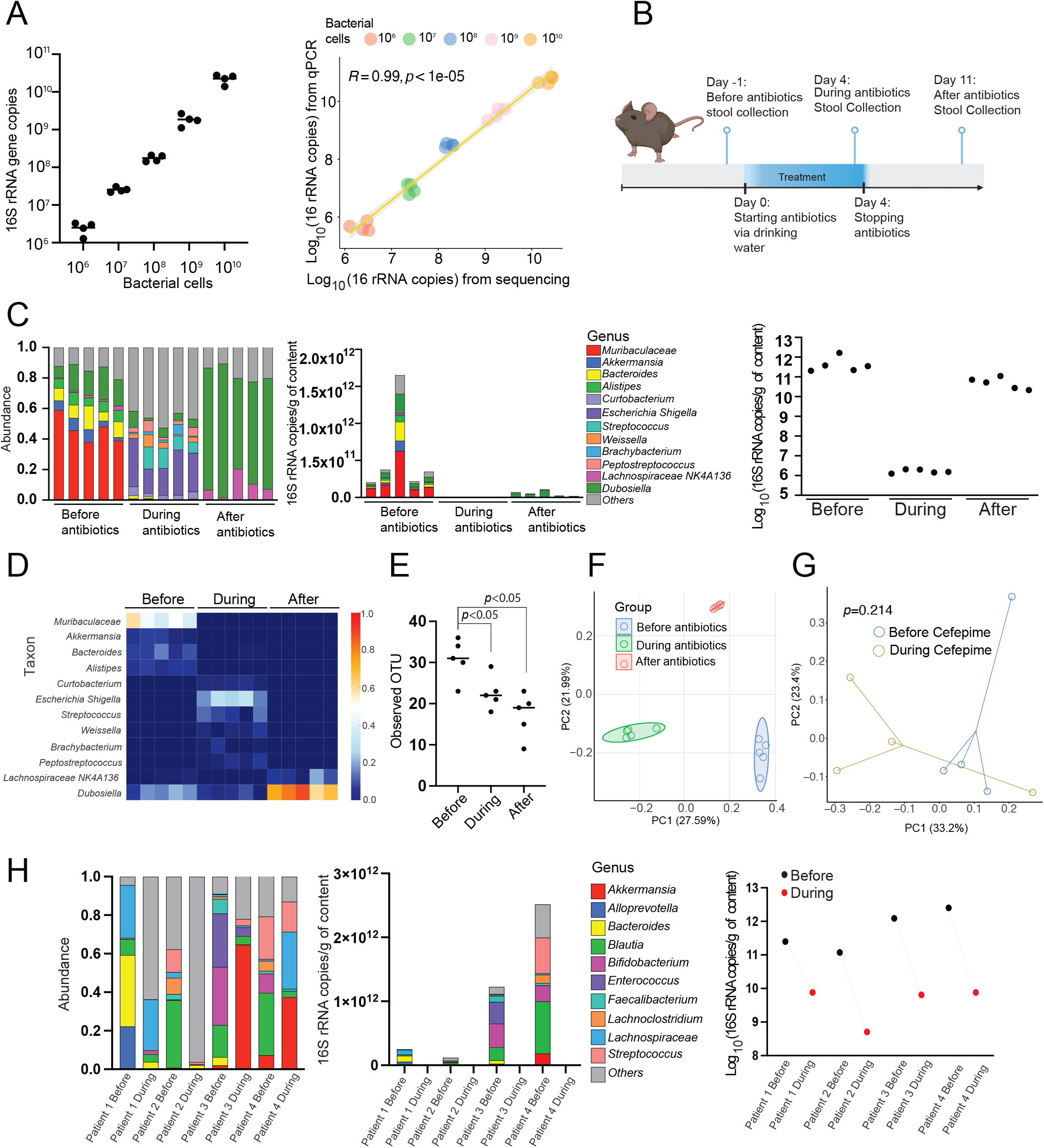
Absolute quantification using BSSI DNA standards compared to traditional quantification methods. (A) Characterization of the absolute quantification methods using *Escherichia coli DH5*_α_ cells. The number of 16S rDNA copies was determined by the spike-in standards (left). Pearson’s correlation analysis compared 16S qPCR to the BSSI method for bacterial count experiments. Pearson R =0.99, P value <1e-05 (right). X and y-axis were logarithm of 16S copy number generated by spike-in standards methods and16S qPCR, respectively. (B) The antibiotic treatment and timeline. (C) The stacked bar charts depict the bacterial composition in mouse samples before, during, and after discontinuing antibiotic treatment. The charts display the relative abundances of bacterial genera (left) and the absolute abundance of 16S rRNA copies per gram of fecal content (middle) of bacterial genera. The absolute abundance of the logarithm of 16S copy number quantification by BSSI spike-in standards (right). (D) A heatmap shows the significant changes in genera between before, during and after decontaminating antibiotic treatment. (E) Alpha diversity, measured as the number of observed OTUs, was quantified in fecal samples. Statistical significance was determined by the Mann-Whitney U test. (F) Principal coordinates analysis (PCoA) using the weighted UniFrac distance for a mouse cohort before, during and after antibiotic treatment, assessed by PERMANOVA testing. (G) PCoA analysis using the weighted UniFrac distance for a patient cohort before and during cefepime treatment, assessed by PERMANOVA testing. (H) The stacked bar charts illustrate changes in bacterial composition in patient samples before and during antibiotic treatment, presenting relative abundances at the genus level (left) and absolute abundance at the genus level (middle). The absolute abundance of the logarithm of 16S copy number quantification by BSSI spike-in standards (right).

### Assessing spike in references for absolute quantification in antibiotic-treated samples

One of the most impactful and potentially disruptive factors influencing the microbiota and outcomes of patients is the administration of antibiotics.^19,29^ Antibiotic treatment is also a common tool for manipulating the microbiota in animal models. We assessed the performance of balanced spike-in standards across a range of murine fecal samples with varying bacterial biomass secondary to antibiotic treatment, evaluating samples collected before and after broad-spectrum antibiotic administration, as well as during the microbiome recovery phase following cessation of antibiotic treatment. A cocktail of vancomycin, ampicillin, and enrofloxacin was administered in the drinking water of mice for four days. Mouse fecal samples were collected immediately before, on the last day of treatment, and a week after stopping antibiotic treatment (Fig. 2B). From the 16S deep sequencing results, we were also able to assess relative abundance of taxonomical composition of the samples, depicted using stacked bar graphs (Fig 2C left panel and 2D). Decontamination of antibiotic treatment reduced the relative abundances of several genera including *Muribaculaceae, Akkermansia, Bacteroides*, and *Alistipes* while enriching the abundances *Curtobacterium, Escherichia-Shigella, Streptococcus, Weissella, Brachybacterium*, and *Peptostreptococcus*. After discontinuing the antibiotic treatment, there was an increase in *Dubosiella* genus and *Lachnospiraceae_NK4A136* genus (Fig. 2C left panel and 2D). We also performed the 16S rRNA copies per gram of fecal pellets for genera in a stacked bar chart (Fig. 2C, middle) and a log scale of total 16S rRNA copies per gram of fecal pellets using the BSSI method (Fig. 2C, right). During antibiotic treatment, a substantial reduction in the mean 16S copy number was observed, decreasing from 5.6E+11 to 1.7E+6 copies/gram compared to before antibiotic treatment. After ceasing antibiotics for 7 days, the mean total of 16S rRNA gene copies increased back to 5.7E+10 copies/gram (Fig. 2C, right). Alpha diversity in the observed OTU analysis was significantly reduced in the during and after antibiotics groups compared to the before antibiotics group (Fig. 2E). A distinct difference in microbial composition was noted using weighted UniFrac analysis between the ‘before’ and ‘during’ groups (P=0.009), as well as between the ‘before’ and ‘after antibiotics’ groups (P=0.006) using PERMANOVA testing (Fig. 2F). These results suggest that balanced synthetic spike-in standards can quantify 16S rRNA copies for genera before, during, and after antibiotic treatment phases and provide important contextual information regarding absolute bacterial densities.

Additionally, spike-in quantification was applied to fecal samples from four patients undergoing stem cell transplantation collected before and during treatment with cefepime. During-treatment samples from patients were collected at least four days after beginning treatment. Patient characteristics are summarized in Table 3. Using principal coordinates analysis (PCoA) with PERMANOVA testing of weighted UniFrac beta diversity measures (P=0.214), microbial composition between the before and during antibiotics groups was not drastically different (Fig. 2G). A stacked bar plot illustrated the shifts in the relative and absolute abundance of microbial composition before and during cefepime treatment for four patients (Fig. 2H, left and middle). The patient samples were quite heterogeneous in their baseline composition. We applied the spike-in method to quantify the total 16S rRNA gene copies before and during cefepime treatment; patients received more than four days of cefepime treatment showed a trend of decrease in mean of total 16S rRNA gene copies significantly reduced to below 7.62E+09 (Fig. 2H, right). Interestingly, the relative abundances of the *Akkermansia* genus increased in patient 3 and patient 4 during cefepime treatment. However, the absolute abundances of the *Akkermansia* genus showed a reduction in patient 3 and patient 4 during cefepime treatment compared to before treatment (Fig. S3A). Similarly, the relative abundances of the *Bacteroides* genus slightly increased in patient 2 during cefepime treatment. However, the absolute abundance of the *Bacteroides* genus showed a decrease during cefepime treatment compared to before treatment (Fig. S3A). These results demonstrate that antibiotic treatment led to reduced gut microbiome densities compared to before treatment. The relative abundances of the changed taxa, however, do not accurately indicate whether the taxa were expanding or shrinking during the treatment. The spike-in standards enable the quantification of changes in microbial densities per gram of stool content in real time, providing a better understanding of the dynamic changes within microbiome.

### Assessment of spike-in standards on sequencing run quality

Amplicon libraries of low diversity, such as the 16S rRNA gene, are commonly supplemented with a high-diversity library such as PhiX to reduce the problem of homogeneous signals during sequencing on the Illumina MiSeq platform^30^. Nonetheless, this strategy has a drawback: it reduces the overall number of reads without additional benefits. To evaluate whether a BSSI approach can improve the diversity and sequencing quality of 16S amplicon sequencing, we first created a mixture of imbalanced spike-in standard mixtures (SP1-pCR, SP2-pCR, SP5-pCR, and SP6-pCR plasmids (Table 1 and 2). The example of design of imbalanced nucleotides in the SP5 and SP6 spike-ins are highlighted in red in Fig. 1A. We also generated model samples composed of only two murine-derived isolates, *Akkermansia muciniphila* (MDA-JAX AM001) and *Bacteroides thetaiotaomicron* (MDA-JAX BT001) (Fig. 3A), to experimentally create a low-diversity sequencing run. Four imbalanced spike-in standards were pre-mixed in a equalmolar ratio and added to samples during lysis step, after which they were constructed into a library alongside the model samples and then conducted 2 × 150 bp paired-end MiSeq runs. We assessed the effect of the imbalanced spike-in standards on several parameters, including overall run quality, cluster density, successful merging percentages, and quality scores (Q scores). A quality score predicts the probability of an error in base calling. A high Q score indicates that a base call is more reliable and has a lower chance of being incorrect.^31^ The accuracy of a sequencing platform is traditionally assessed by the percentage of Q30 scores which correspond to a base call accuracy of 99.9%, meaning there is a 1 in 1000 chance of an incorrect base call. To generally enhance data quality for low-diversity samples, PhiX DNA was added to the final library at approximately 6%-7% for runs. The cluster densities were 1059.6 ± 37 for the run without spike-ins and 809.3 ± 7 for the run with imbalanced spike-ins. The pass filter rates were 89.66% ± 1.5% for run without spike-ins, compared to 87.57% ± 0.54% for the run with. The run without spike-ins had a successful merging percentage of 98.64%, whereas it was 95.5% for the run with imbalanced spike-ins (Table 5). Moreover, we compared the percentages of bases equal to and exceeding Q39, Q38, or Q30 in both read 1 and read 2. The Miseq runs with imbalanced spike-ins exhibited a significant decrease in the percentages of Q39, Q38, and Q30 in the second read, compared to the runs without spike-in standards (Fig. 3B).

**Figure 3.**
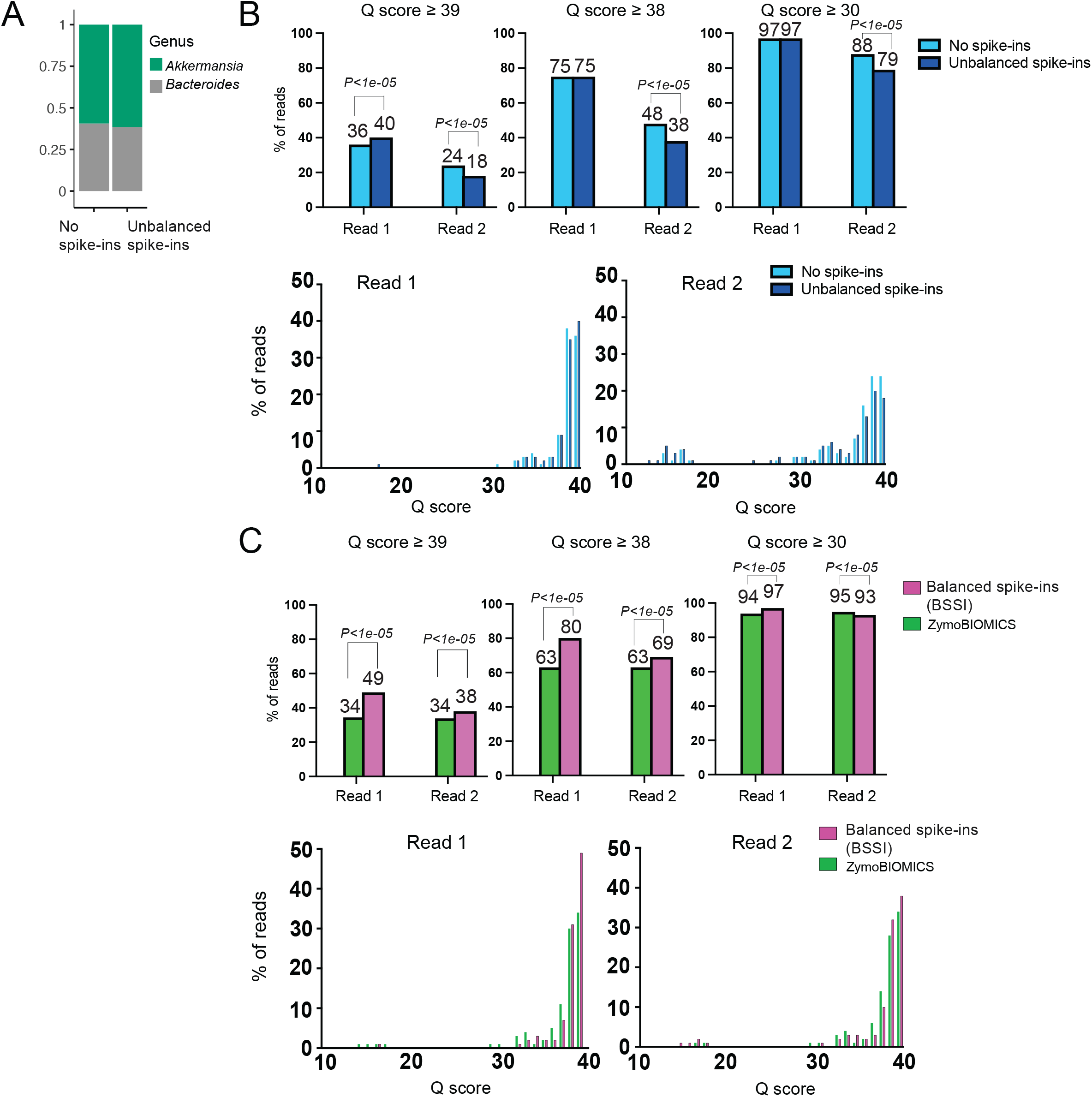
Evaluation of impact of imbalanced and balanced spike-in standards on paired-end sequencing run quality. (A) Evaluation of imbalanced spike-in standards on sequencing run quality using a model sample. Relative abundance of *Akkermansia muciniphila* and *Bacteroides thetaiotaomicron* from runs with and without adding imbalanced spike-in DNA standards in the lysis step. (B) Q scores ≥39, ≥38, and ≥30 for reads 1 and 2 from runs with and without imbalanced spike-in standards are shown in the plots (upper). We performed pairwise proportional tests on Q scores ≥39, ≥38, and ≥30 between runs with and without the imbalanced spike-in for reads 1 and 2. The spiking in of the imbalanced standards mix significantly reduced the Q scores ≥39, ≥38, and ≥30 on read 2, with p-values <1e-05. Similarly, the imbalanced standards mix lowered the Q scores ≥39 on read 1 compared to the runs without standards, with p-values also <1e-05. The histograms overlay the distribution of the percentage of reads per Q score for Read 1 and Read 2, with (dark blue) and without (light blue) imbalanced spike-in standards. The x-axis represents the Q score, and the y-axis represents the percentage of reads (lower). (C) Q scores ≥39, ≥38, and ≥30 for reads 1 and 2 from runs with the addition of either BSSI standards or commercial ZymoBIOMICS Spike-in Control II that are displayed in the plot (upper). A pairwise proportional test was performed between runs with and without the addition of either spike-in standards or commercial spike-in bacteria in the antibiotic-treated mouse fecal DNA. The addition of the balanced standards mixes significantly increased the Q scores ≥39, ≥38, and ≥30 for read 1 compared to the runs without the spike-in mix, with p-values <1e-05. Similarly, the BSSI standards mix increased the Q scores ≥39 and ≥38 for read 2 compared to the runs with ZymoBIOMICS standards, with p-values also <1e-05. However, the standards mix slightly reduced the Q scores ≥30 for read 2 compared to the runs with ZymoBIOMICS standards. The histograms overlay ZymoBIOMICS standards (green) and BSSI spike-in standards (pink), showing the distribution of the percentage of reads per Q score for Read 1 and Read 2, with BSSI standards and ZymoBIOMICS spike-in standards (lower).

We then further evaluated the potential impact of balanced spike-in standards on Illumina sequencing quality metrics, particularly for low diversity and low biomass samples following antibiotic decontamination. We compared our balanced spike-in standards with the commercially available ZymoBIOMICS Spike-in Control II standards. This ZymoBIOMICS standards is designed for cell number measurements from low bacterial loads in NGS-based analysis and consists of three bacterial strains: *Truepera radiovictrix, Imtechella halotolerans*, and *Allobacillus halotolerans*, mixed at different cell counts of 2.0 × 10^5^, 3.0 × 10^4^, and 7.0 × 10^3^ per 20 μL of solution. The ZymoBIOMICS standards did not contain evenly distributed nucleotides present in every nucleotide. We asked if introducing these to low-density samples could potentially result in lower quality performance due to the low nucleotide diversity from imbalanced nucleotide representation. We compared our BSSI standards with the commercially available ZymoBIOMICS standards and evaluated fecal samples collected from CD11c-Cre^+^ Stat3^f/f^ mice. This mouse model, which has a deletion of STAT3 from dendritic cells, lacks STAT3’s anti-inflammatory function and underwent an antibiotic cocktail treatment as part of an immune checkpoint inhibitor-induced autoinflammatory colitis model.^14^ Genomic DNA was extracted from fecal pellets collected from these mice after five days of antibiotic treatment to produce low biomass samples. To achieve more than 75% of the sequences, balanced spike-in standards and ZymoBIOMICS standards were used to mimic standards that occupy the majority of the reads in low biomass samples. We extracted bacterial DNA from ZymoBIOMICS standards. ZymoBIOMICS standards DNA and our BSSI DNA were co-amplified low biomass DNA from mice to assess the sequencing quality performance of spike-in strategies. We conducted 2 × 150 bp paired-end sequencing runs without adding PhiX DNA and assessed their effects on various parameters, including overall run quality and Q scores (Table 5). The total abundances of ZymoBIOMICS standards consisting of three bacteria, Truepera radiovictrix (26.6%), Imtechella halotolerans (44%), and Allobacillus halotolerans (4.5%), occupied approximately 75% of the reads, whereas BSSI occupied 77.5%. We controlled for cluster density and passing filter rates to ensure the sequencing runs had similar conditions. The cluster density was 678.9 ± 7 for runs with four balanced spike-ins and 683.85 ± 13 for runs with ZymoBIOMICS standards. The passing filter rates were 86.55% ± 0.51% for runs with spike-ins and 89.09% ± 6.78% for runs with ZymoBIOMICS standards (Table 5). Notably, our BSSI approach resulted in improved quality scores compared to ZymoBIOMICS standards, with significantly better Q39 and Q38 percentages for both read 1 and read 2 (Fig. 3C). Overall, these results suggest that the addition of balanced spike-in standards allows for absolute quantification without adversely affecting run quality parameters and that the balanced nucleotide approach could offer enhancements in quality scores relative to imbalanced spike-in approaches.

## Discussion

Incorporating spike-in standards as internal references in high-throughput sequencing can significantly enhance the rapid quantification of microbial densities across numerous samples. In our study, we introduced the reference DNA in the lysis buffer before DNA extraction, allowing the spike-in standards to reflect extraction efficiency. Given that amplicon libraries can be of low diversity, we opted for synthetic DNA standards to achieve balanced nucleotide representation at each position, which we hypothesized could result in superior sequencing quality scores. Our approach demonstrated significant higher Q39 and Q38 percentages for both read 1 and read 2 compared to commercial ZymoBIOMICS spike-in standards. Additionally, BSSI method eliminated the need for additional PhiX to maintain diversity while achieving high Q scores without occupying some of the read depth by PhiX. This underscores the critical role of spike-in design, particularly for low biomass samples, by providing quantitative information on total bacterial loads and economically benefiting the sequencing run by reducing the amount of PhiX required.

We evaluated different normalization methods commonly used in statistics; however, they have not been widely tested for generating standard curves describing the relationship between relative abundances of gene counts and absolute gene densities. A previous study used spiking known amounts of exogenous bacteria into samples to appropriately normalize the total microbial loads, a process called spike-in-based bacterial abundance calibration (SCML)^10^. They applied log_2_-transformed read counts of known concentrations of spiked-in bacteria for the calibration of ratios of absolute abundance. This method was shown to reduce variation and accurately calculates ratios of abolute OTU abundaces of samples. Another study generated standard curves for absolute quantification to convert read counts to absolute copy numbers, based on aggregated dose-response curves obtained from a mix of spike-in standards with various concentrations, which were fitted using negative binomial Poisson generalized linear models (GLM) with a fixed slope of 1.^32^ We found that using logit normalization for the relative abundances of gene counts produced a better R-squared value, as would be anticipated given that relative abundances are bounded from 0 to 1 (Fig. S1A). Alternative approaches, including arcsine or anscombe normalization, produced curves with slopes that decrease and flatten at the extremes of gene densities evaluated (Fig. S1B and S1C). This indicates that logit normalization could be a more accurate model for describing the relationship between absolute and relative abundances in the setting of spike-in quantification.

Our method can be potentially extended to outpatient collection preservatives by incorporating a spike-in standard mix into commercially available preservatives. After collection, the stool and preservative mixtures can undergo the 16S rRNA sequencing workflow for absolute quantification per gram of fecal samples without additional quantitative PCR. Our method has shown the capability to detect total bacterial densities across a large dynamic range (Fig 2A, 2C and 2H). Absolute abundances generated by BSSI standards can be correlated with those obtained from 16S qPCR by Pearson correlation analysis (Fig. 2A). Absolute quantification of microbial densities is crucial for accurately understanding associations with disease in samples from pre-clinical models or patients. In our mouse model, after ceasing antibiotics for 7 days, the mean 16S copy number increased to 5.7E+10 (Fig. 2C), which is only a 10-fold reduction compared to the mean 16S copy number before antibiotic treatment. However, alpha diversity in the observed OTU analysis was still significantly lower after ceasing antibiotics for 7 days compared to before antibiotic treatment (Fig. 2E). We quantified the absolute abundance for each genus and found that *Dubosiella* and *Lachnospiraceae_NK4A136* were not significantly expanded during the microbial recovery phase after ceasing antibiotics compared to before treatment. However, the relative abundance of these genera significantly increased during the microbial recovery phase compared to before antibiotic treatment (Fig. S3B). These data suggest that it would be beneficial to have quantitative microbiome profiling for better understanding the mechanism in microbiome reserach.

Emerging evidence suggests that microbial load analysis based on 16S amplicon sequencing is needed for biomarker identification.^33^ Spike-in standard concentrations, however, may need further adjustment, particularly with very low bacterial density samples, such as tumor and FFPE samples. Previous studies have demonstrated that droplet digital PCR (ddPCR) can accurately measure species-specific bacterial DNA in blood during bloodstream infections.^34^ For samples with very low biomass, it is important to prevent potentially over-occupying reads with standards, which results in low-resolution quantification of true sample-derived sequences. It could be beneficial to use ddPCR to qualify, determine, and adjust the spike-in concentrations. By adding the optimal spike-in mix before extraction, we can not only measure absolute bacterial loads but also prevent potentially over-occupying reads with standards, resulting in low-resolution quantification of true sample-derived sequences.

## Conclusions

We have developed synthetic standards for absolute quantification method that can be readily incorporated into standard 16S deep sequencing workflows using BSSI standards. Spike-in read counts are identified and computationally removed from sequencing counts derived from actual samples, allowing usual downstream relative abundance quantification workflows. This allows concurrent quantification of total bacterial loads in addition to relative abundances in microbiome samples. We implemented our method across a range of bacterial counts, spanning five orders of magnitude which provide highly accurate and reliable results. Absolute abundance generated by BSSI standards can be correlated with 16S quantitation PCR method by Pearson correlation analysis. We found that this strategy also performed well in different biomass samples such as fecal samples collected from mice and patients before and after antibiotic treatment. Our mixture of BSSI standards includes four different plasmids designed to produce equal nucleotide representation at each position, which we found resulted in improved run quality parameters compared to commercial spike-in methods. To conclude, introducing balanced synthetic spike-in standards is a powerful tool that can be adapted to various types of samples by providing absolute microbial abundance quantification.

## Declarations

### Acknowledgments

We thank Dr. Stephanie Watowich for the CD11c-Cre+ Stat3f/f mice (Stat3Δ/Δ).

## Author Contributions

CCC, YJ, TH, MO, IF, LM, and IG contributed to sample processing. CCC and RRJ contributed to conceptualization and methodology. CCC and IG contributed to clinical sampling and clinical data collection. CCC, LH, YJ, MJ, EH, TH, MO, DP, IG, IF, LM, RP, CP, and RRJ conducted stool and antibiotics studies and analyzed/interpreted the data. All authors approved the manuscript.

## Availability of data and materials

All data generated in this study are openly available.

---

Supplementary Figure 1. The formula for the BSSI standard curve was calculated using the known 16S rRNA gene copies on a log_10_ scale as the x-axis, related to relative abundances of the pCR4-*Blautia* plasmid from 16S V4 deep sequencing after applying the normalization methods indicated as the y-axis. (A) The BSSI standard curve is based on the logit normalization of the pCR4-*Blautia* plasmid and the log_10_ of the known 16S gene copies of the pCR4-*Blautia* plasmid. (B) The BSSI standard curve is based on the arcsine normalization of the pCR4-*Blautia* plasmid and the log_10_ of the known 16S gene copies of the pCR4-*Blautia* plasmid. (C) The BSSI standard curve is based on the anscombe normalization of the pCR4-*Blautia* plasmid and the log_10_ of the known 16S gene copies of the pCR4-*Blautia* plasmid.

Supplementary Figure 2. Agarose gel analysis of PCR amplification for four BSSI standards, SP1, SP2, SP3 and SP4. 2% agarose gel showed that all four BSSI standards has around 390 base pairs amplicon, including adaptors and inserts.

Supplementary Figure 3. (A) The relative abundance and absolute 16S copy number quantification by BSSI standard methods for genera *Akkermansia* and *Bacteroides* in a patient cohort before and after treatment with cefepime. (B) Relative abundances of the genera *Dubosiella* and *Lachnospiraceae_NK4A136* in mouse samples before, during, and after discontinuing antibiotic treatment, quantified by 16S rRNA gene sequencing. The bar indicates the mean of all values. Statistical significance was determined by the Mann-Whitney U test.

Table 1. DNA sequences of synthetic spike-ins SP1, SP2, SP3 and SP4.

Table 2. DNA sequences of synthetic spike-ins SP5 and SP6.

Table 3. Characteristics of the study patients.

Table 4. Standard curve for BSSI quantification.

Table 5. Overall run quality on Miseq runs.

## Reference

1 Rooks, M. G. & Garrett, W. S. Gut microbiota, metabolites and host immunity. Nat Rev Immunol 16, 341–352 (2016). 10.1038/nri.2016.42

2 Chen, Y. et al. Oncogenic collagen I homotrimers from cancer cells bind to alpha3beta1 integrin and impact tumor microbiome and immunity to promote pancreatic cancer. Cancer Cell (2022). 10.1016/j.ccell.2022.06.011

3 Wei, M. et al. High-Throughput Absolute Quantification Sequencing Revealed Osteoporosis-Related Gut Microbiota Alterations in Han Chinese Elderly. Front Cell Infect Microbiol 11, 630372 (2021). 10.3389/fcimb.2021.630372

4 Boer, C. G. et al. Intestinal microbiome composition and its relation to joint pain and inflammation. Nat Commun 10, 4881 (2019). 10.1038/s41467-019-12873-4

5 Vandeputte, D. et al. Quantitative microbiome profiling links gut community variation to microbial load. Nature 551, 507–511 (2017). 10.1038/nature24460

6 De Tomassi, A. et al. Combining 16S Sequencing and qPCR Quantification Reveals Staphylococcus aureus Driven Bacterial Overgrowth in the Skin of Severe Atopic Dermatitis Patients. Biomolecules 13 (2023). 10.3390/biom13071030

7 Wang, X., Howe, S., Deng, F. & Zhao, J. Current Applications of Absolute Bacterial Quantification in Microbiome Studies and Decision-Making Regarding Different Biological Questions. Microorganisms 9 (2021). 10.3390/microorganisms9091797

8 Ubeda, C. et al. Vancomycin-resistant Enterococcus domination of intestinal microbiota is enabled by antibiotic treatment in mice and precedes bloodstream invasion in humans. J Clin Invest 120, 4332–4341 (2010). 10.1172/JCI43918

9 Yang, L., Lou, J., Wang, H., Wu, L. & Xu, J. Use of an improved high-throughput absolute abundance quantification method to characterize soil bacterial community and dynamics. Sci Total Environ 633, 360–371 (2018). 10.1016/j.scitotenv.2018.03.201

10 Stammler, F. et al. Adjusting microbiome profiles for differences in microbial load by spike-in bacteria. Microbiome 4, 28 (2016). 10.1186/s40168-016-0175-0

11 Celis, A. I. et al. Optimization of the 16S rRNA sequencing analysis pipeline for studying in vitro communities of gut commensals. iScience 25, 103907 (2022). 10.1016/j.isci.2022.103907

12 Oldham, A. L. & Duncan, K. E. Similar gene estimates from circular and linear standards in quantitative PCR analyses using the prokaryotic 16S rRNA gene as a model. PLoS One 7, e51931 (2012). 10.1371/journal.pone.0051931

13 Zaramela, L. S., Tjuanta, M., Moyne, O., Neal, M. & Zengler, K. synDNA-a Synthetic DNA Spike-in Method for Absolute Quantification of Shotgun Metagenomic Sequencing. mSystems 7, e0044722 (2022). 10.1128/msystems.00447-22

14 Zhou, Y. et al. Intestinal toxicity to CTLA-4 blockade driven by IL-6 and myeloid infiltration. J Exp Med 220 (2023). 10.1084/jem.20221333

15 Takeda, K. et al. Stat3 activation is responsible for IL-6-dependent T cell proliferation through preventing apoptosis: generation and characterization of T cell-specific Stat3-deficient mice. J Immunol 161, 4652–4660 (1998).

16 Caton, M. L., Smith-Raska, M. R. & Reizis, B. Notch-RBP-J signaling controls the homeostasis of CD8-dendritic cells in the spleen. J Exp Med 204, 1653–1664 (2007). 10.1084/jem.20062648

17 Wang, Y. et al. Author Correction: Fecal microbiota transplantation for refractory immune checkpoint inhibitor-associated colitis. Nat Med 25, 188 (2019). 10.1038/s41591-018-0305-2

18 Schwabkey, Z. I. et al. Diet-derived metabolites and mucus link the gut microbiome to fever after cytotoxic cancer treatment. Sci Transl Med 14, eabo3445 (2022). 10.1126/scitranslmed.abo3445

19 Hayase, E. et al. Mucus-degrading Bacteroides link carbapenems to aggravated graft-versus-host disease. Cell 185, 3705–3719 e3714 (2022). 10.1016/j.cell.2022.09.007

20 Rognes, T., Flouri, T., Nichols, B., Quince, C. & Mahe, F. VSEARCH: a versatile open source tool for metagenomics. PeerJ 4, e2584 (2016). 10.7717/peerj.2584

21 Edgar, R. C. UNOISE2: improved error-correction for Illumina 16S and ITS amplicon sequencing. bioRxiv (2016).

22 Schloss, P. D. et al. Introducing mothur: open-source, platform-independent, community-supported software for describing and comparing microbial communities. Appl Environ Microbiol 75, 7537–7541 (2009). 10.1128/AEM.01541-09

23 Quast, C. et al. The SILVA ribosomal RNA gene database project: improved data processing and web-based tools. Nucleic Acids Res 41, D590–596 (2013). 10.1093/nar/gks1219

24 Caballero, S. et al. Cooperating Commensals Restore Colonization Resistance to Vancomycin-Resistant Enterococcus faecium. Cell Host Microbe 21, 592–602 e594 (2017). 10.1016/j.chom.2017.04.002

25 Yang, Y. W. et al. Use of 16S rRNA Gene-Targeted Group-Specific Primers for Real-Time PCR Analysis of Predominant Bacteria in Mouse Feces. Appl Environ Microbiol 81, 6749–6756 (2015). 10.1128/AEM.01906-15

26 Parada, A. E., Needham, D. M. & Fuhrman, J. A. Every base matters: assessing small subunit rRNA primers for marine microbiomes with mock communities, time series and global field samples. Environ Microbiol 18, 1403–1414 (2016). 10.1111/1462-2920.13023

27 Apprill, A., McNally, S., Parsons, R. & Weber, L. Minor revision to V4 region SSU rRNA 806R gene primer greatly increases detection of SAR11 bacterioplankton. Aquat Microb Ecol 75, 129–137 (2015). 10.3354/ame01753

28 Gilbert, J. A., Jansson, J. K. & Knight, R. The Earth Microbiome project: successes and aspirations. BMC Biol 12, 69 (2014). 10.1186/s12915-014-0069-1

29 Stein-Thoeringer, C. K. et al. A non-antibiotic-disrupted gut microbiome is associated with clinical responses to CD19-CAR-T cell cancer immunotherapy. Nat Med 29, 906–916 (2023). 10.1038/s41591-023-02234-6

30 Naik, T., Sharda, M. C P. L., Virbhadra, K. & Pandit, A. High-quality single amplicon sequencing method for illumina MiSeq platform using pool of ‘N’ (0-10) spacer-linked target specific primers without PhiX spike-in. BMC Genomics 24, 141 (2023). 10.1186/s12864-023-09233-4

31 Ewing, B. & Green, P. Base-calling of automated sequencer traces using phred. II. Error probabilities. Genome Res 8, 186–194 (1998).

32 Tourlousse, D. M. et al. Synthetic spike-in standards for high-throughput 16S rRNA gene amplicon sequencing. Nucleic Acids Res 45, e23 (2017). 10.1093/nar/gkw984

33 Tito, R. Y. et al. Microbiome confounders and quantitative profiling challenge predicted microbial targets in colorectal cancer development. Nat Med 30, 1339–1348 (2024). 10.1038/s41591-024-02963-2

34 Ziegler, I., Lindstrom, S., Kallgren, M., Stralin, K. & Molling, P. 16S rDNA droplet digital PCR for monitoring bacterial DNAemia in bloodstream infections. PLoS One 14, e0224656 (2019). 10.1371/journal.pone.0224656

