## Supplementary figures and images for "Accurate quantitation of 16S gene copies in low biomass samples post-antibiotic treatment through deep sequencing with a balanced nucleotide synthetic spike-in approach"

### Supplemental Figure 1

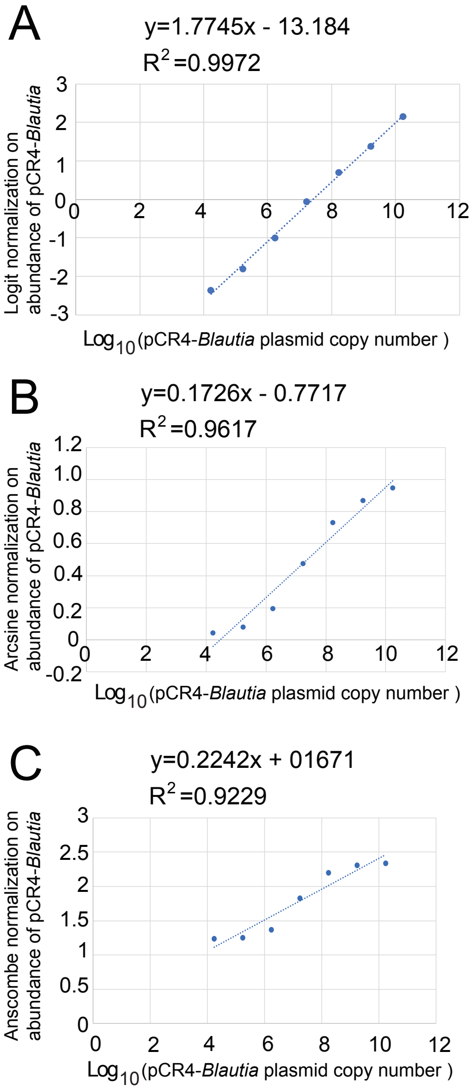

### Supplemental Figure 2

Figure S2

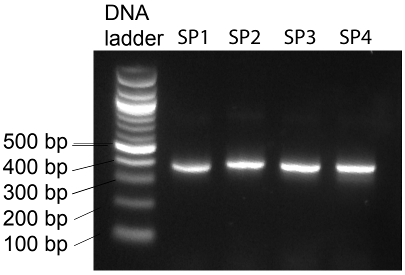

### Supplemental Figure 3

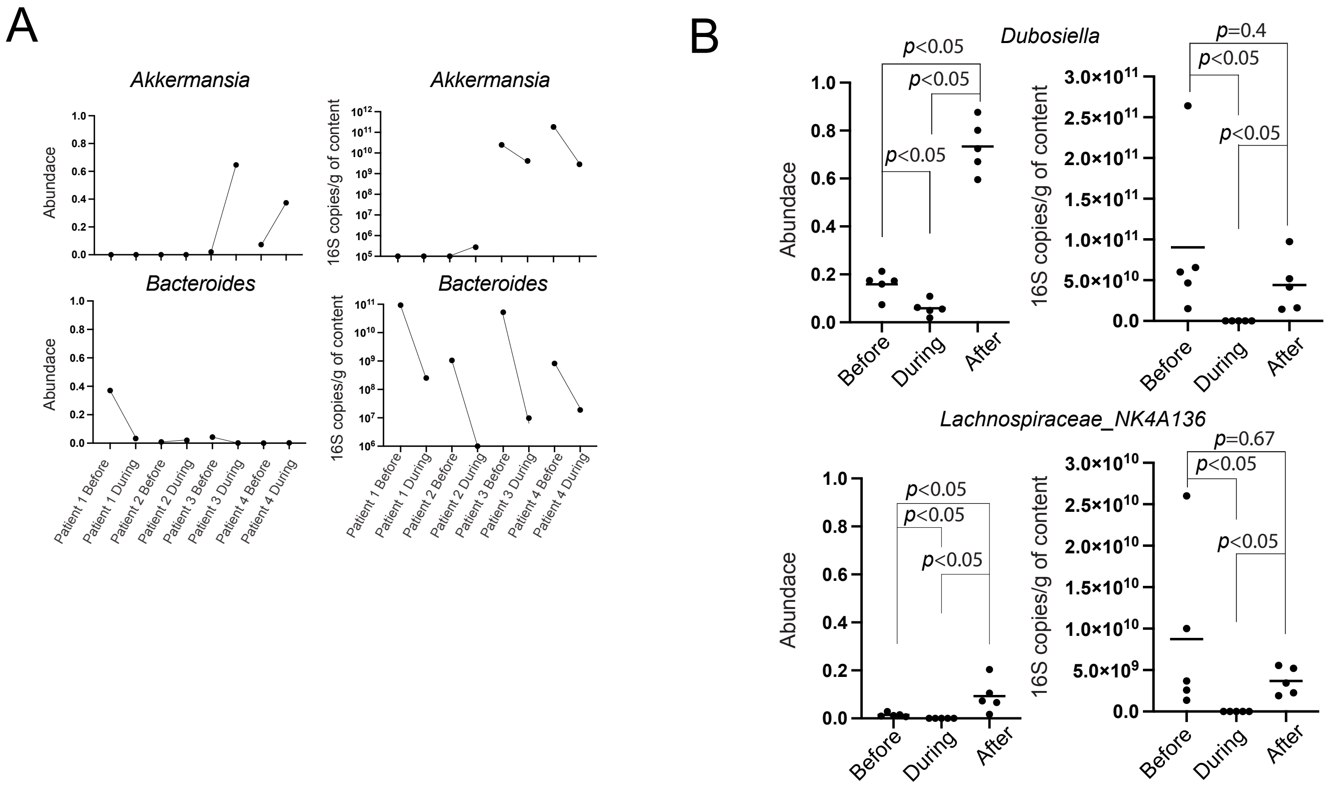
